# Self-sustained rhythmic behavior of *Synechocystis* PCC 6803 under continuous light conditions in the absence of light-dark entrainment

**DOI:** 10.1101/2023.09.26.559469

**Authors:** Lutz C. Berwanger, Nikolaus Thumm, Rahil Gholamipoor, Anika Wiegard, Jeannine Schlebusch, Markus Kollmann, Ilka M. Axmann

## Abstract

Circadian clocks regulate biological activities, providing organisms a fitness advantage under diurnal changing conditions by allowing them to anticipate and adapt to recurring external changes. In recent years attention was drawn to the entrainment by intracellular cycles. Photosynthetic Cyanobacteria coordinate their gene expression, metabolism, and other activities in a circadian fashion. Solely, three proteins, KaiA, KaiB, and KaiC, constitute the well-studied circadian clock of the cyanobacterial model, *Synechococcus elongatus* PCC 7942. It remained inconclusive for a long time whether *Synechocystis* sp. PCC 6803, an important organism for biotechnological applications, can also maintain circadian rhythms under continuous illumination. Using an approach, which does not require genetic modification, we investigated the growth behavior of *Synechocystis* via non-invasive online backscattering measurement and verified all three criteria for true circadian oscillators: temperature compensation, entrainment by external stimuli, and a self-sustained freerunning period of about 24 hours. Since manipulation of the circadian clock (*Synechocystis* Δ*kaiA1B1C1*) led to a significant reduction in glycogen content, disruption of glycogen synthesis (*Synechocystis* Δ*glgC*) entirely inhibited glycogen formation and both mutants lost oscillations, we hypothesize that the oscillations reflect glycogen metabolism.

**Significance Statement:** Monitoring circadian rhythms in cyanobacteria usually requires genetically modified reporter strains or intensive sampling for downstream analysis. Even for the main cyanobacterial model *Synechocystis* sp. PCC 6803 it was debated for years to which extent undamped circadian oscillations are really present until a suitable reporter strain was developed. We applied online backscatter measurements as an alternative readout to monitor circadian oscillations in cyanobacteria. In *Synechocystis* the temperature-compensated *kaiA1B1C1*-driven 24 h metabolic oscillations did not require light-dark entrainment, highlighting the relevance of the clock for the carbon metabolism even under continuous light, an aspect which should be considered for industrial set-ups. Our method opens the possibility to extend circadian analysis to non-GMO and monitor metabolic rhythmicity during high-density cultivation.

## Introduction

From a theoretical perspective, an oscillation is a repetitive or periodic change of a measure around a central value or between two or more different states. In a biological context, a common form of oscillation is the circadian rhythm, which represents a self-sustained oscillation with a period of approximately 24 hours. Natural biological activities follow circadian patterns that allow organisms to adapt to daily environmental changes due to the Earth’s rotation. This adaptation can be seen in a range of organisms, from humans to cyanobacteria (1). Three criteria define a circadian oscillator: the first attribute is the presence of a self-sustaining oscillation with a period of 24 hours. The second characteristic is the ability to synchronize (entrain) the internal oscillator with external rhythmic stimuli (*Zeitgeber*). The third criterion is that the period length of the endogenous oscillation is not significantly affected by ambient temperature and remains constant over a physiologically relevant temperature range (2–4).

Cyanobacteria, a monophyletic group of photoautotrophic prokaryotes, have been used for decades to study fundamental processes such as photosynthesis and gene regulation, and are attractive hosts for biotechnological production (5, 6). They are well-suited model-organisms for studying circadian rhythms and the clock’s connection to metabolism (7–9). In cyanobacteria, diurnal changes in metabolism and transcription occur and are associated with the output of the cyanobacterial circadian clock (10, 11). A timing system that enables them to predict changes in light before sunrise or sunset and to adjust the expression of certain genes, such as for photosynthesis, is beneficial (12, 13). This protein system, composed of the Kai proteins − KaiA, KaiB, and KaiC − forms the central oscillator for endogenous timing in cyanobacteria (4, 14, 15). In *Synechococcus elongatus* PCC 7942 (*Synechococcus*), circadian regulation has been shown for many physiological phenomena, including gene expression and chromosome compaction (16, 17). KaiC phosphorylation oscillates around the diurnal course and, as an oscillator, rhythmically regulates gene expression via an output apparatus that controls cyanobacterial physiology (18, 19). While in *Synechococcus*, the circadian clock has been shown to be the dominant factor in controlling gene expression, the clock of *Synechocystis* sp. PCC 6803 (*Synechocystis*) appears to control transcription to a much lesser extent (20, 21). In contrast to *Synechococcus*, multiple kai gene copies exist in *Synechocystis*: two *kaiA (kaiA1, kaiA3)*, three *kaiB* (*kaiB1 - B3*), and three *kaiC* homologs (*kaiC1-C3*) (22–24). It is most likely that KaiA1B1C1 represents the *bona fide* oscillator (21, 23, 25–28). Transcriptomic analyses of a Δ*kaiA1B1C1* deletion mutant showed altered expression of genes related to metabolic processes, such as photosynthesis, respiration, and carbon metabolism, as well as altered translational and transcriptional regulation (26). A very recent promoter study showed that circadian oscillations are diminished after *kaiA1B1C1* deletion (28). This suggests that KaiA1B1C1 provides the coordination of cellular timing, probably in a cross-talk with KaiB3 and KaiC3 (28, 29). In the model organism *Synechococcus*, KaiC interacts either directly or indirectly with circadian output components like the regulator of phycobilisome associated A (RpaA), histidine kinase *Synechococcus* adaptive sensor A (SasA), and CikA (18, 19). Until now, the output signaling pathway of the putative central KaiA1B1C1 oscillator in *Synechocystis* has not been completely clarified. Orthologs for both SasA and RpaA are present, and deletion of either encoding gene leads to growth deficits in the light-dark (LD) cycle, especially under mixotrophic conditions (21). *In vivo* interactions with the SasA ortholog (Hik8, sll0750) have only been shown for KaiC1 and not for KaiC3 (8).

Glycogen is the main storage compound in *Synechocystis* and *Synechococcus* and is anabolized during the day and catabolized at night, and is known to be regulated by the clock proteins in *Synechococcus* (30–34). Diurnal metabolism begins with the transfer of carbon flux from the oxidative pentose phosphate pathway (OPPP) to the Calvin-Benson-Bassham cycle (CBBC) and is governed by products of photosynthetic light reactions (35–37). During the day, the CBBC captures CO_2_ and diverts the excess carbon into glycogen stores (38). ADP-glucose pyrophosphorylase (glucose-1-phosphate adenyltransferase, GlgC) − a regulatory enzyme in the anabolism of glycogen (31, 39) − is critical for the biosynthesis of glycogen.

Although *Synechocystis* expresses *kai* gene orthologs and clearly displays diurnal patterns of photosynthesis, respiration as well as diurnal glycogen synthesis and degradation (10, 30, 35, 38, 40), circadian regulation is not well understood yet and the literature is partially conflicting. Furthermore, genome-scale transcription rhythms were not maintained or rapidly attenuated under continuous light conditions (41). Interestingly, the transcription-translation regulatory loop that depends on the KaiC-dependent expression of *kaiBC* does not appear to be present in *Synechocystis* (22).

The coupling of luciferase to a clock-controlled endogenous promoter revealed a functional circadian clock for both *Synechococcus* and *Synechocystis*. The rhythms of the resulting bioluminescence indicate more rapid damping of oscillations in *Synechocystis* than in *Synechococcus* and displayed only low amplitude in *Synechocystis* (28, 42–46). However, very recently true circadian and high amplitude rhythms of promoter activity were elegantly confirmed using a super strong heterogenous promoter (28). Kucho et al. 2005 (20) demonstrated circadian oscillations of less than 9% of genes in continuous light (LL) after stimulation by a single pulse of 12 h of darkness, while in another study, gene expression levels in LL or darkness even suggest diurnal rhythmicity rather than circadian oscillation (41). On the contrary, sustained circadian rhythms under LL and LD were described in 2015 for *Synechocystis* in a photobioreactor (47). Shortly after, diurnal oscillating behavior of approx. 40% of the genes in *Synechocystis* were revealed, which were involved in several cellular processes (10). Altogether, the previous studies imply that a circadian control is present in *Synechocystis*, but the activity of endogenous promoters might not cycle with such a high amplitude as in *Synechococcus*, while the metabolism and growth might still be under circadian control. The majority of studies reporting self-sustained rhythms have in common that they used non-invasive systems, where the cultures were not disturbed by taking samples for down-stream analysis. We, therefore, set out to establish a non-invasive method that can detect circadian oscillations without the need of detecting gene expression rhythms.

We used online backscatter measurements to demonstrate that *Synechocystis* WT and *Synechococcus* WT display self-sustained oscillations with a ∼24 h period in the backscatter signal under constant light conditions. Our method allows non-invasive live monitoring of non-GMO batch cultures under well-defined conditions, such as light, atmosphere, humidity, and temperature. The rhythms did not even require entrainment by environmental cycles, but cells were synchronized by a single nutrition upshift. The backscatter signal oscillations were temperature-compensated and therefore circadian in nature. Our results show that the oscillations were directly governed by the circadian clock since the oscillation pattern was lost in the *kaiA1B1C1* deletion mutant. We conclude that glycogen metabolism is impaired in this mutant as it almost does not accumulate glycogen (∼1% CDW). Both observations suggest a direct correlation between the oscillatory backscatter signal and glycogen metabolism. Thus, our research provides valuable insights into the metabolic pathways and circadian rhythm of *Synechocystis*, a cyanobacterial strain, widely utilized in biotechnology and chronobiology research.

## Results

### *Synechocystis* displays 24 h backscatter oscillations in a non-invasive system

To investigate whether growth occurs in a circadian rhythm in the cyanobacterium *Synechocystis* we studied growth behavior based on non-invasive online backscatter measurements. After an initial dilution in fresh medium, *Synechocystis* WT batch cultures displayed steady growth with an average doubling time of ∼7 days (based on backscatter signal). However, albeit being grown under constant conditions, we observed daily changes in the (overall increasing) raw backscatter signals (Fig. 1A). The same behavior was observed for *Synechococcus* WT (Fig. 1D), which was grown as a control. Backscatter signals had varying noise levels due to the shaking of the cultures. We, therefore, performed a time-series data analysis (II Materials and Methods “Data analysis and visualization”) of the backscatter signal, which revealed stable oscillations (Fig. 1A, 1D). A subsequent wavelet transformation showed that recurrent oscillations of the signal occurred throughout the experimental duration − the oscillations clustered at a period of 10 h to 45 h for both WT strains (Fig. 1B, Fig. 1E). To assess the significance of the oscillations in the wavelet spectrum, we compared the results to the gaussian distributed red noise spectrum. The periods of the oscillations that were above the 5% confidence interval had significantly different durations of 24.85 hours ± 0.32 for *Synechocystis* WT (Fig. 1C) and 25.65 hours ± 0.47 for *Synechococcus* WT (Fig. 1F). To rule out the possibility that the backscatter signals were artifacts due to backscatter measurement bias, we performed the experiments investigating *Synechocystis* WT (n=11) and *Synechococcus* WT (n=5) under standard conditions (80 μmol_photons_ m^-2^ s^-1^, 30°C, 150 rpm, 0,5% CO_2_) multiple times.

**Figure 1:**
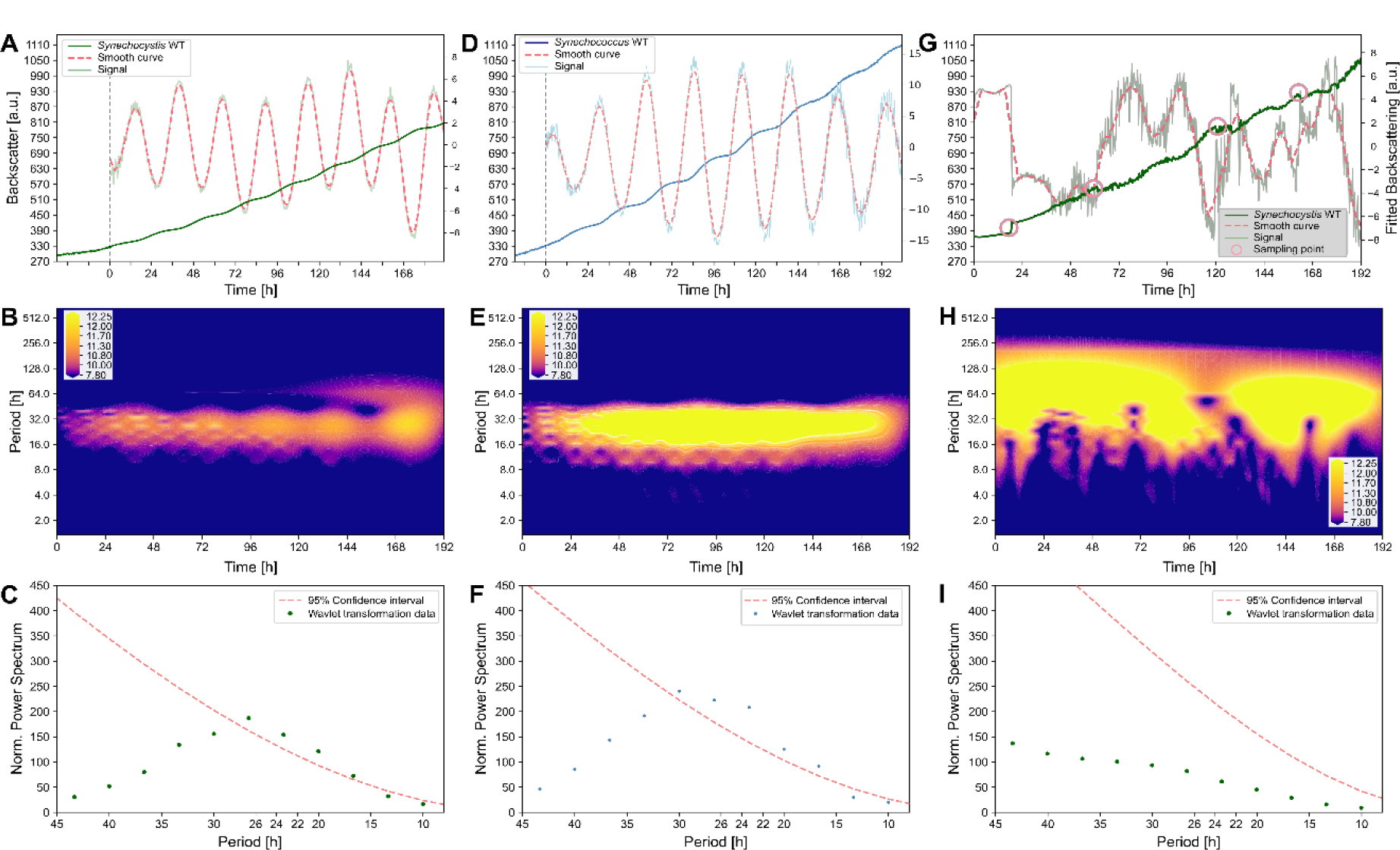
Online backscatter signal analysis of *Synechocystis* sp. PCC 6803, *Synechococcus elongatus* PCC 7942, and invasive sampling. **A**: Representative raw Backscatter signal of *Synechocystis* WT (dark green, left y-axis) batch cultures under constant standard conditions, plotted from the start of the experiment, defined by the initial dilution of the culture. The time, when the backscatter signal reached the threshold of 330 backscatter units, was defined as time 0 for further data processing. “Signal” (light green, right y-axis) displays the denoised signal. “Smooth curve” (dashed red line, right y-axis) displays the predominant oscillation. **B**: Heatmap displaying the periods over time calculated by wavelet trans-formation of the extracted signal. Color scales indicate low to high coefficients **C:** Null-Hypothesis test for the periods obtained by wavelet transformation (green dots). Respective periods (x-scale) are plotted against the power spectrum from the wavelet transformation (y-scale). Gaussian noise-based curve displayed as “95% confidence interval” (dashed red line). **D**: Representative backscatter signal of *Synechococcus* WT (blue) batch cultures treated and displayed as in A. **E+F**: Downstream process of signal analysis as in B, C for *Synechococcus* WT (D). **G**: Representative backscatter signal after invasive sampling of *Synechocystis* WT batch culture (green) under constant conditions. Red circles indicate sampling time points. For sampling, the culture was taken out of the light incubator and returned after taking out 100 μl culture under sterile conditions. **H+I**: Downstream process of signal analysis as in B, C for the invasive sampling run (G).

Sampling for time series is challenging since the continuous sampling of the same batch culture leads to a disturbance of the backscatter signal and might therefore interrupt the ∼24 h rhythm. We tested the effect of taking samples and indeed observed disruption of the backscatter signal (Fig. 1G, n=2). Taking only four samples over the course of eight days disturbed the rhythms so tremendously that no period could be determined anymore (Fig. 1H, I). Thus, our further setup is non-invasive, and we do not interrupt the signal by sampling or other interventions.

### Oscillations of the backscatter signal are temperature compensated

Observing a self-sustained ∼24 h period for *Synechocystis* WT motivated us to investigate whether the oscillations might be of circadian nature. To test for temperature compensation, we repeated the experiment at 25°C, referring to Aoki, Kondo, and Ishiura (44). Periods did not differ significantly between the temperatures and displayed a Q_10_ value of 1.01 (Fig. 2B), being close to previous reports for *Synechocystis* (Q_10_ = 1.08 - 1.1 (28, 42, 47) and characteristic for circadian clocks (48). The observed temperature compensation was confirmed by a single experiment at 35°C (Fig2B). Notably, *Synechococcus* WT backscatter signal amplitude oscillated less at 25°C compared to 30°C and 35°C (Fig. 2D). In addition, the backscatter amplitude of *Synechocystis* WT (∼5/-5 a.u., Fig. 2A) was only one-third as high as the amplitude of *Synechococcus* WT (∼15/-15 a.u., Fig. 2C).

**Figure 2:**
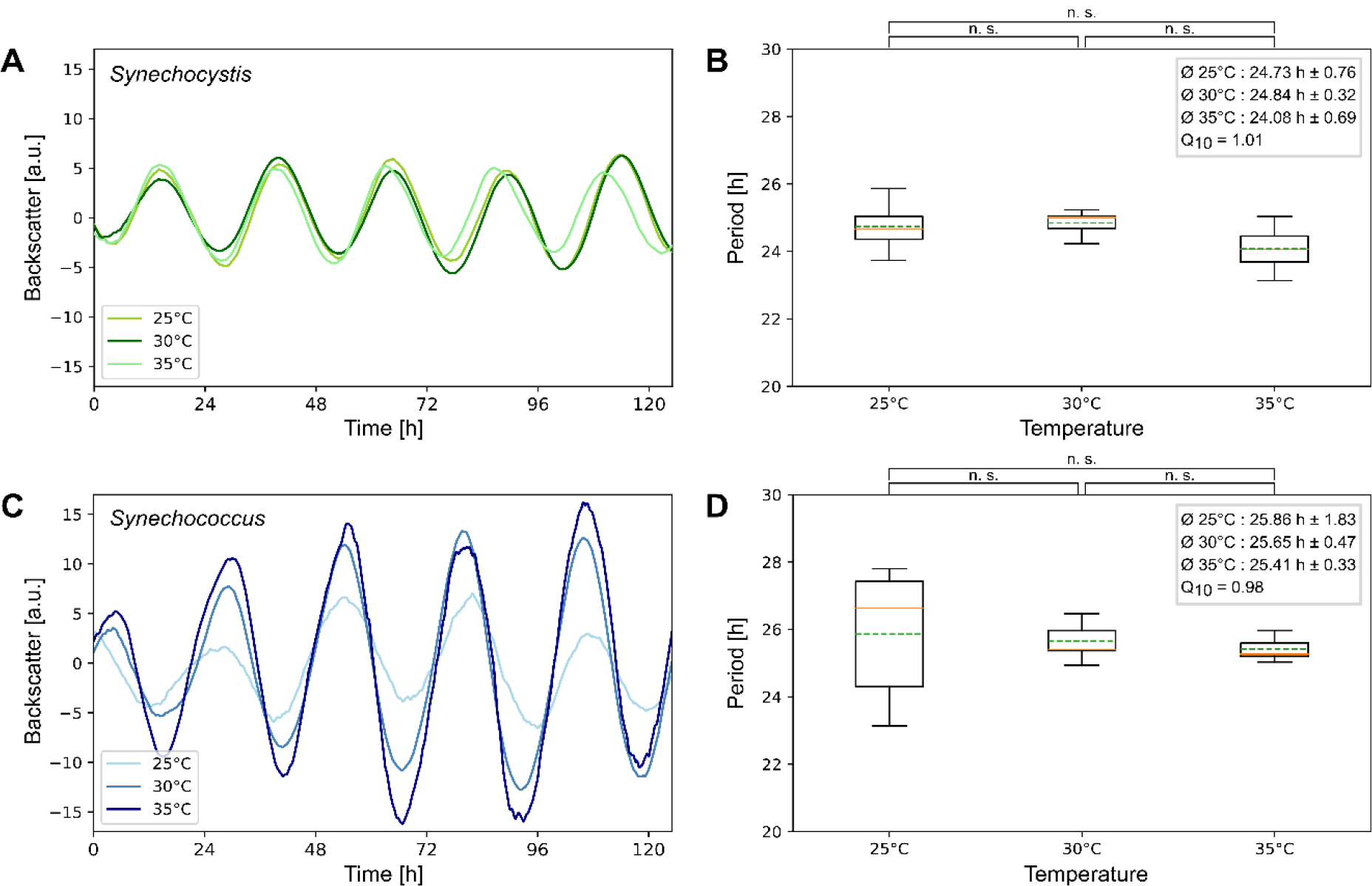
Temperature compensation of the backscatter oscillations. **A**: Overlay of representative backscatter signals for *Synechocystis* WT (green) batch cultures for different temperatures (as indicated in legend). Other conditions remain constant (80 μmol_photons_ m^-2^ s^-1^, 150 rpm, 0,5% CO_2_). **B**: Boxplots displaying the periods determined at the respective temperatures. Peaks were determined using the *“SciPy*.*signal*.*find_peaks*” function. The means (dashed green lines) and the medians (solid orange lines) of the periods are indicated. Temperature compensation was confirmed by 1.) independent t-tests for the three temperatures (p-values ≧0.05). Additionally, Q_10_ values of 1 are within the accepted range (88). The overall average for all temperatures of *Synechocystis* WT was 24.61 h ± 0.66, and *Synechococcus* WT was 25.64 h ± 0.95. **C, D**: Corresponding graphs for *Synechococcus* WT (blue). **D**: t-test p-values ≧0.05, Q_10 values_=1.01 (derived from 25°C and 30°C, were validated with a singular 35°C measurement).

### Backscatter oscillations can be entrained

The above described oscillations were observed in continuous light without entrainment by light-dark cycles. We speculated that the initial dilution of the cultures served as a nutritional upshift and could synchronize the culture. To test this hypothesis, we set up batch cultures as we did before, but we initiated them 12 hours apart. One culture was diluted with fresh BG-11 medium to an OD_750nm_ of 0.8 (light green in Fig. 3A), and a second batch culture was diluted 12 h later from a second pre-culture (dark green in Fig. 3A). After processing and overlaying the backscatter data of the two cultures we observed a shift of roughly 12 hours (11.80 +/-2.47, n=3) between the peaks of their backscatter oscillation, indicating a successful synchronization (Fig. 3B).

**Figure 3:**
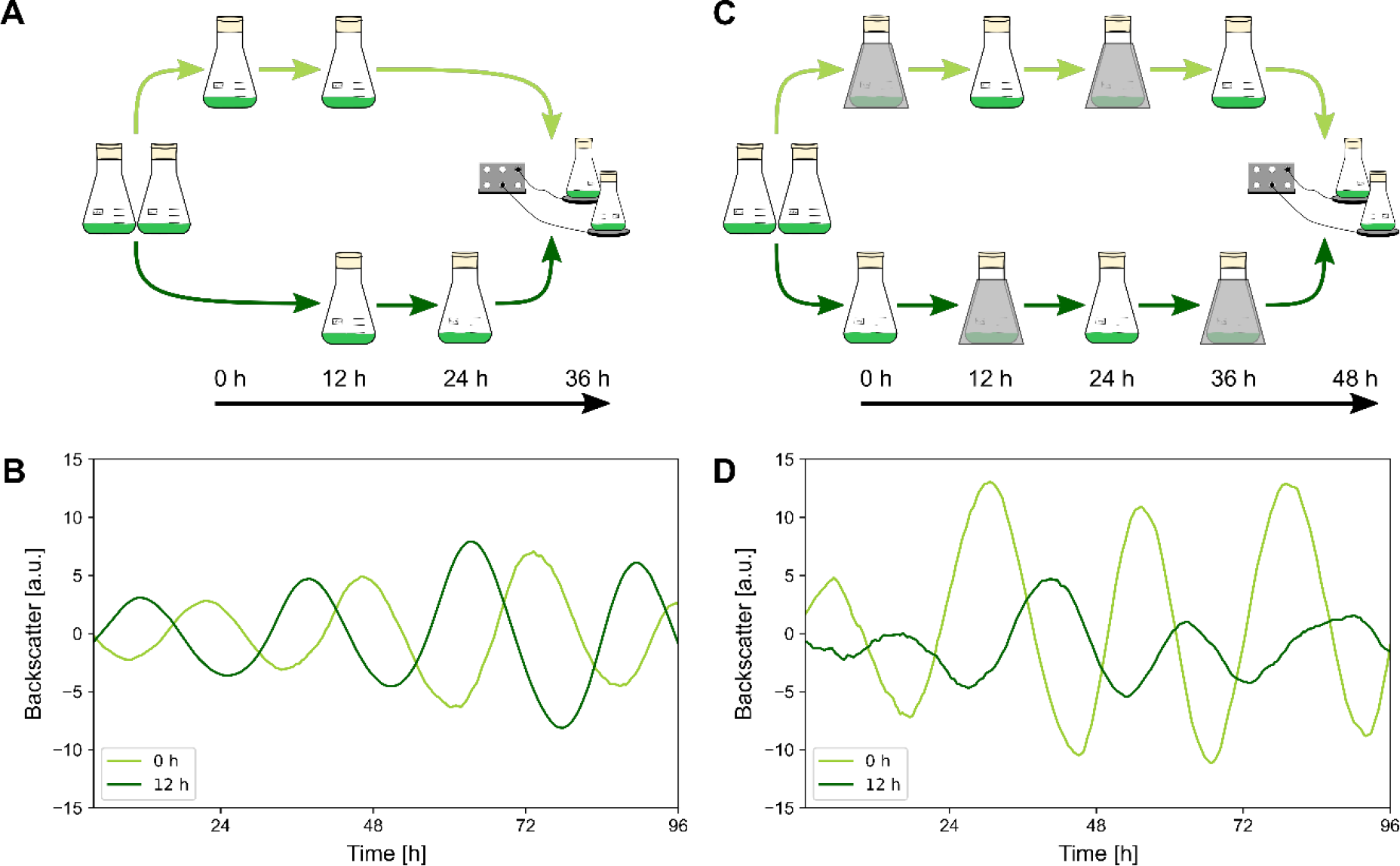
Entrainment of the *Synechocystis* WT oscillations. **A**: Schematic illustration of entrainment by 12 h nutrient shift. **B**: Representative overlay of backscatter signals for *Synechocystis* WT batch cultures separated and entrained by a 12 h nutrient upshift over 48 h. The first culture diluted to an OD_750nm_ of 0.8 at the time point “0 h” (light green), and the second culture diluted to an OD_750nm_ of 0.8 12 h apart (“12 h”, dark green). **C**: Schematic illustration of 12 h LD entrainment over 48 h. **D**: Representative overlay of backscatter signals for *Synechocystis* WT batch cultures separated and entrained by a 12 h LD cycle over 48 h (as indicated in the legend). The first culture (“0 h”, light green) was exposed to LD cycles (12:12 LD, OD_750nm_ of 0.8) over 48 h and the second culture (“12 h”, dark green) corresponding to “0 h” but 12 h apart.

An important criterion for a circadian oscillation is that the intrinsic rhythm can be synchronized to an environmental rhythm. Therefore, we exposed two cultures to 12:12 light-dark pulses after the initial nutrient-upshift. We shifted the light-dark cycles for two cultures by 12 hours to test whether we can introduce a phase delay (Fig. 3C). If the dark pulses were given in phase with the nutrition upshift (light green in Fig. 3D) stable oscillations were measured. If the dark pulses were given 12 hours later (dark green in Fig. 3D), the amplitude of the oscillatory signal decreased and was less smooth in Fig. 3D, “12 h”, but shifted by ∼12 hours (11.23 +/-7.7, n=3). Since both cultures (Fig. 3D, “0 h” and “12 h”) were LD entrained, the disrupted signal may be due to weaker effects on the oscillator compared to nutrition upshift.

### Backscatter oscillations are connected to the glycogen metabolism and are diminished in a *Synechocystis* clock mutant

Given that the observed oscillations fulfilled the criteria of circadian oscillations (period duration of about 24 hours, which can be entrained by external stimuli and remains stable over different temperatures), we sought to test whether oscillations are affected by manipulation of the clock. For this purpose, we created a knock-out mutant (Δ*kaiA1B1C1*) and applied the experimental setting to these mutants as described above. As shown in Figure 4A (n=4), the deletion of the putative KaiA1B1C1 core clock disrupted the oscillating pattern.

**Figure 4:**
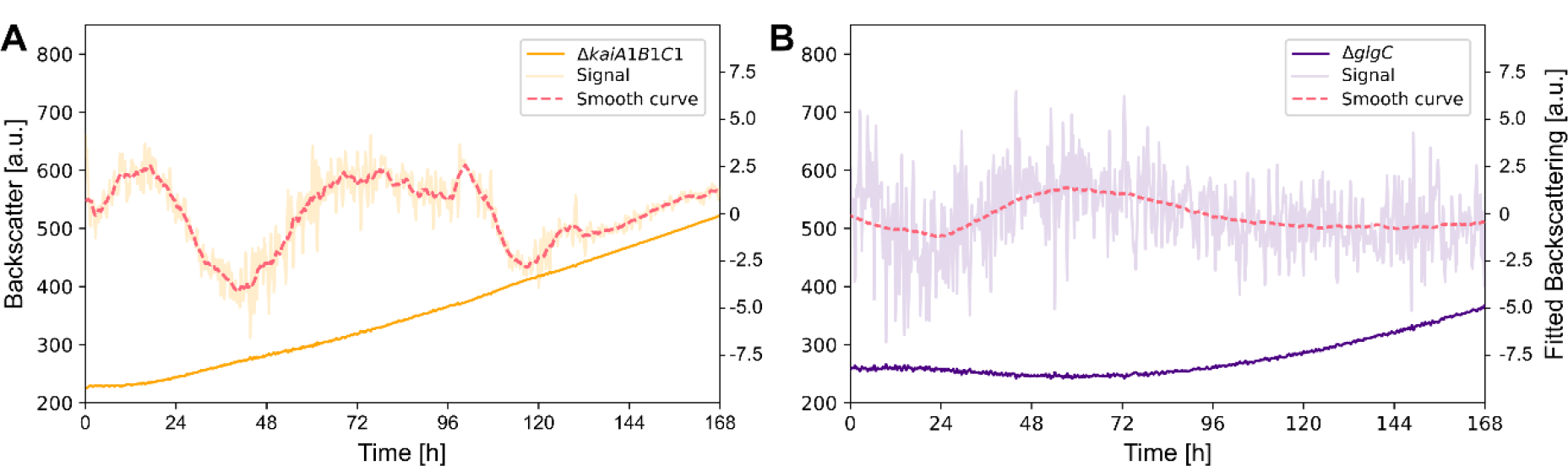
Online backscatter signal analysis of *Synechocystis* Δ*kaiA1B1C1* and Δ*glgC*. **A**: Backscatter signal of *Synechocystis* Δ*kaiA1B1C1* (orange). “Signal” (light orange) displays the denoised signal. “Smooth curve” (dashed red line) displays the predominant oscillation. **B**: Backscatter signal of *Synechocystis* Δ*glgC* (indigo) batch cultures under constant conditions. “Signal” (light indigo) displays the denoised signal. “Smooth curve” (dashed red line) displays the predominant oscillation.

For *Synechococcus* WT, one of the most prominent outputs of the circadian clock is the regulation of glycogen metabolism (34, 49). Glycogen is synthesized during the day and catabolized during the night when light energy is limited (38, 40). Diel glycogen oscillations were also observed for *Synechocystis* (30, 35). We, therefore, aimed to investigate whether the observed backscatter oscillations can be attributed to changing glycogen contents. The *Synechocystis* WT accumulated 16% glycogen per CDW, which fits the observations (Ø = 18,5% per CDW) from Velmurugan and Incharoensakdi in 2018 (50). We created a knockout mutant (*Synechocystis* Δ*glgC*) that is unable to synthesize glycogen by deletion of the *glgC* gene ((51) and Fig. 5B). In fact, no oscillation pattern was seen in Δ*glgC* (Fig. 4B, n=3), confirming that the backscatter signal oscillations are connected to changes in glycogen. A comparison of the two *Synechocystis* deletion mutants and the WT is shown in Fig. 5A. By measuring the glycogen content (Fig. 5B) 86 h to 96 h after nutrition upshift for the three strains (Fig. 5A), we could determine that *Synechocystis* Δ*glgC* cannot synthesize glycogen and that *Synechocystis* Δ*kaiA1B1C1* displayed strongly decreased glycogen content per cell dry weight (∼1% CDW^-1^) implying that the loss in backscatter oscillations might originate from dysregulated glycogen metabolism.

**Figure 5:**
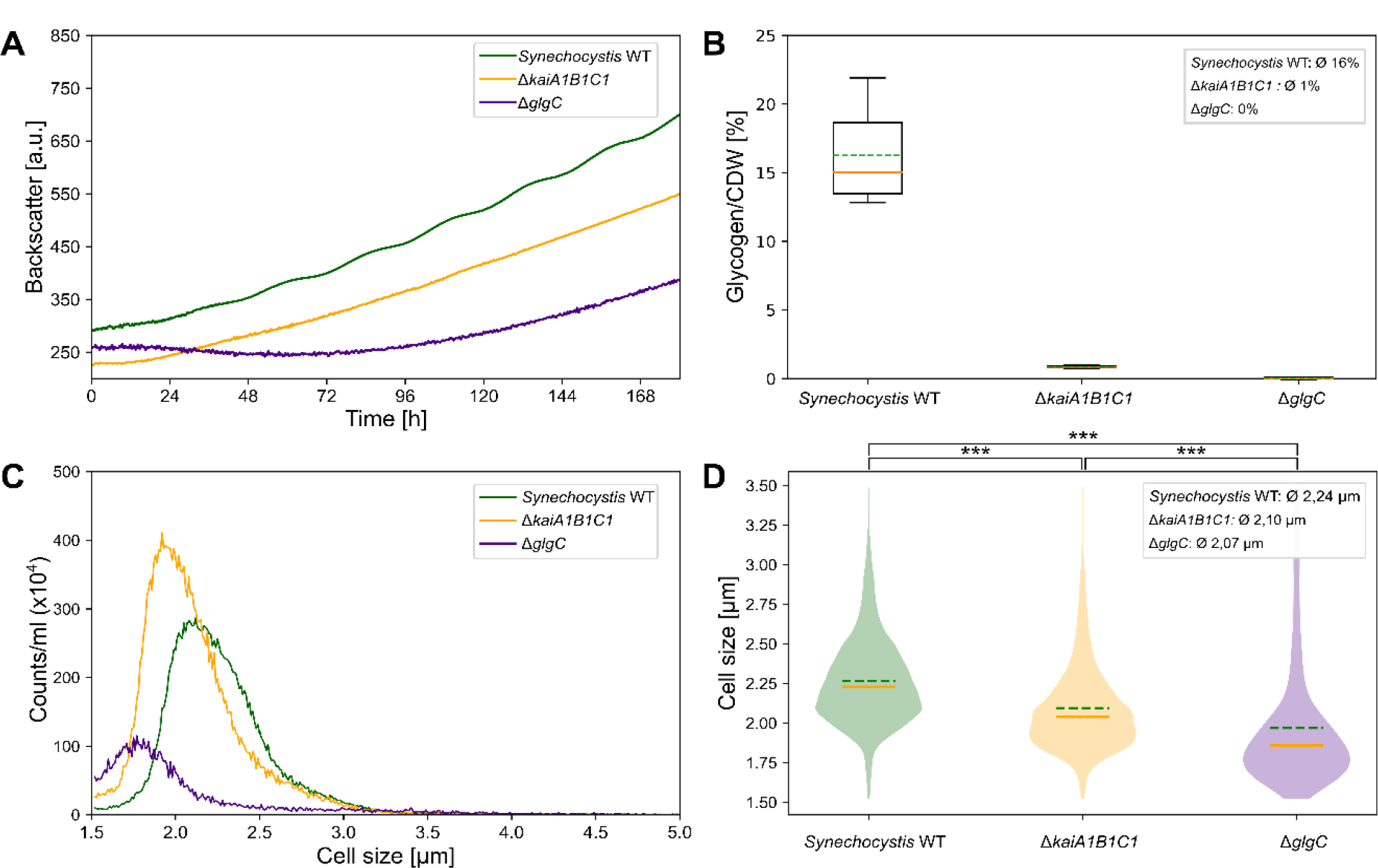
Comparison of physiological analysis of *Synechocystis*. **A**: Representative backscatter signals of *Synechocystis* WT (green), *Synechocystis* Δ*kaiA1B1C1* (orange), and *Synechocystis* Δ*glgC* (indigo). **B**: Boxplots displaying the glycogen content per CDW. The means (dashed green lines) and the medians (solid orange lines) of the Ø content for each strain are indicated. Only values above 0 are shown. 86 h after inoculation Samples were taken over a period of 12 h, every 2 h from WT cultures or every 4 h for the mutant strains, respectively. **C**: Cell size and cell count distribution of *Synechocystis* strains. **D:** Corresponding violin plots of C with the associated density curves of the respective cell sizes. The independent t-tests for mean counts of the three *Synechocystis* strains have the p-values < 0.001. The means (green dashed lines) and the medians (solid orange lines) of the cell size for each strain are indicated. **B-D:** Data of cell analysis study.

We determined the cell count and the distribution of cell sizes using an electric field cell counting system (Fig. 5C). The overall cell counts correlated with the growth curves derived from the backscatter signal (Fig. 5C). In correlation with the glycogen content, the three strains, *Synechocystis* WT, *Synechocystis* Δ*glgC*, and *Synechocystis* Δ*kaiA1B1C1*, showed differences in cell size. *Synechocystis* WT cells displayed a wide size range and were on average larger (mean cell size ∼2.24 μm) than the mutant cells, with decreasing size from *Synechocystis* Δ*kaiA1B1C1* (mean cell size ∼2.10 μm) to *Synechocystis* Δ*glgC* cells (mean cell size ∼2.07 μm). *Synechocystis* Δ*glgC* and *Synechocystis* Δ*kaiA1B1C1* cells showed a narrow peak and, hence a smaller variety of cell sizes compared to the *Synechocystis* WT (Fig. 5C, D). Altogether, our measurements show a correlation between easy-to-monitor backscatter oscillations and the cellular glycogen content, which affects cell size and is regulated by the circadian clock.

## Discussion

### Detection of circadian backscatter oscillations in a non-invasive system

Over decades *Synechococcus* served as a perfect model for chronobiology (28, 52). The attractiveness of cyanobacteria as hosts for bioproduction (53, 54), and the emerging evidence of oscillations in other prokaryotes (55) call for extending the understanding of circadian clocks to further organisms. So far, most studies on cyanobacterial circadian rhythms relied on live monitoring of reporter strains or measuring gene expression rhythms after sampling via labor- and cost-intensive transcriptomic analysis. The case of *Synechocystis* illustrates that establishing a reporter construct requires a lot of optimization. It took 30 years after the first reports of promoter activity rhythms until a very strong sensor was published recently (28, 42, 46). However, gene transcript oscillations also do not necessarily overlap with protein rhythms and metabolic oscillations (56). The here established live-monitoring of the cyanobacterial growth behavior provides a fast alternative for screening cyanobacteria for circadian rhythms.

Backscatter measurement is the method of choice for high-cell density cultivation since it is accurate at OD ≧1 to 70, while optical density measured as absorbance is only efficient in the OD range of 0.3 to 0.8 (57, 58). Here, backscatter measurements facilitated monitoring oscillations in *Synechococcus* WT and *Synechocystis* WT over several days without disturbing the culture. Since the periods of oscillations in the two strains at 30°C differed by ∼1 h in the exact same setup (Fig. 1B, D), we can exclude that the periodic fluctuations are due to external cues (e.g., opening of the light incubator). In agreement with a recent study, which reported the dependence of promoter activity rhythms on *kaiA1B1C1*, the backscatter oscillations were driven by the core oscillator (28).

### Entrainment by nutrition up-shift and light-dark cycles

Most studies on circadian activity in cyanobacteria applied a light-dark regime for initial synchronization of the culture before investigating oscillations in free-running conditions, because the fitness advantage lies in an adjustment of the internal rhythms to the cycling environment. Alternatively, temperature-rhythms have been applied (59). Our results suggest that a single nutrient upshift can synchronize oscillations in the batch culture, although we cannot rule out the possibility that the initial dilution of the culture may also act as a light pulse. We were also able to replicate a classical entrainment by two 12-hour LD pulses, which was shown to work for *Synechococcus* (42) and *Synechocystis* before (47), but we observed differences in the amplitudes after shifting the dark pulse by 12 hours. The lowered amplitude might arise from giving the dark pulse when the cells were in a circadian phase which is less responsive to the stimulus, since phase-dependent resetting of the clock - a general phenomenon of circadian rhythms (60) − has been demonstrated for *Synechocystis* before (28, 45). In addition, we hypothesize that nutrition upshift may be a stronger entrainment cue for *Synechocystis* than darkness. Different impacts of external stimuli regarding the entrainment of the circadian clock among different species (e.g., *Neurospora crassa*) have been observed previously (61). Food uptake was shown to be an entrainment signal (62–64) and e.g. sets up the characteristic rhythms of glycogen metabolism in the liver of rats (65) and humans (66, 67). Similarly, an engineered *Synechococcus* strain could be synchronized by rhythmic supply of glucose. Sugar feeding blocked the effect of resetting the clock by a dark pulse, even in the absence of light (62). However, 24h rhythms without LD-entrainment have been observed not only for plants, microalgae, and cyanobacteria in bioreactor-setups (6, 68, 69) but also for non-photosynthetic prokaryotes like *E. coli* (55).

### Glycogen metabolism in *Synechocystis*

The circadian system in *Synechococcus* regulates cellular physiology by adjusting to environmental fluctuations that affect metabolic rhythms (9, 62, 70). This is demonstrated by the observation that the glycogen content in *Synechococcus* wild-type cells oscillates with a period of 24.7 ± 0.13 hours and is controlled by the clock output components SasA and RpaA (8, 11, 34). A knockout of the *kaiBC* genes in *Synechococcus* resulted in the loss of glycogen oscillation (34). In *Synechocystis*, the interaction between KaiA1B1C1 and the SasA ortholog Hik8 and RpaA, respectively, interferes with glycogen metabolism (8, 21, 71, 72). This KaiAB1C1-SasA-RpaA system influences switching the metabolism from photoautotrophy to the utilization of internal carbon reserves (73). The deletion of *kaiA1B1C1* in *Synechocystis* reduced the glycogen content and led to smaller cell sizes in comparison to the wild type, indicating impaired glycogen metabolism. Our hypothesis is that due to the low glycogen content in *Synechocystis* Δ*kaiA1B1C1* and the inability of *Synechocystis* Δ*glgC* to synthesize glycogen, it is likely that other sugar structures are formed from the absorbed CO2 (51, 74, 75). The lack of glycogen storage in *Synechocystis* Δ*kaiA1B1C1* may impair light/dark transitions, as the circadian clock’s main role is to adjust the cell’s physiological state for the upcoming night environment (20).

Glycogen content interferes with backscatter signals due to its light-scattering properties (76, 77), which can bias optical density measurements and potentially obscure oscillation patterns in *Synechocystis* Δ*glgC* mutants. The loss of pattern may result from the inability to detect glycogen fluctuations in the backscatter signal rather than a disruption of circadian rhythms. Similarly, 24-hour rhythms in *Synechocystis* under LL disappeared after manipulation of topoisomerase expression, which increased glycogen content and optical density without affecting cell numbers (6).

### Conclusion/Outlook

In this study, we detected true circadian rhythms with a free-running period of 24.61 h ± 0.66 in *Synechocystis*. Our data indicate that KaiA1B1C1 represents the *bona fide* oscillator in *Synechocystis* and thereby contributes to resolving a long-standing debate about circadian oscillations in *Synechocystis*. We propose that the observed oscillations are linked to the glycogen content of the cell and that the KaiA1B1C1 system in *Synechocystis* regulates the metabolism of this central storage compound. The fact that oscillations could be synchronized by the initial culture dilution implies that synchronized glycogen oscillations might always be present in simple batch cultures.

*Synechocystis* has several Kai homologs and the homologs *kaiB3* and *KaiC3* were proven to be relevant for circadian promoter activity (28). The exact role of these additional clock components also in the context of glycogen metabolism remains to be clarified. Live monitoring of cyanobacteria holds immense potential for biotechnological applications. Diverse strategies and tools have been developed to manipulate carbon flow in cyanobacteria, and glycogen metabolism has been generally recognized as a promising target (74, 78–80). The monitoring method allows researchers to identify optimal time frames for inducing heterologous protein production, synthesizing valuable products by correlating with glycogen content, as recently demonstrated for the circadian clock (81). It also facilitates the optimization of growth conditions for cyanobacteria and provides opportunities for the development of novel biotechnological processes.

## Materials and Methods

### Strains and growth conditions

*Synechocystis* sp. PCC 6803 and *Synechococcus elongatus* PCC 7942 strains were always grown at 30°C, 0,5% CO_2_, 75% humidity, 150 rpm, with constant light of 80 μmol_photons_ m^-2^ s^-1^, in a Multitron Infors HT^®^ Incubator, if not stated otherwise. Cultures were cultivated in BG11 (82) supplemented with a final concentration of 10 mM TES (TES PUFFERAN® ≥99%, Carl Roth) to a final pH of 8 with KOH. Pre-cultures were grown in 100 ml BG11 in 250 ml Erlenmeyer flasks and inoculated from prepared BG11 agar plates. The mutant strains *Synechocystis* Δ*kaiA1B1C1* and *Synechocystis* Δ*glgC* were supplemented with kanamycin (25 μg ml^-1^).

### DNA isolation

For DNA isolation, 0.2% glucose was added to the culture the evening before DNA isolation. 50 ml culture were centrifuged at 6,000 g for 10 min, 4°C, and washed two times in 10 ml TE buffer (10 mM Tris/HCl, 1 mM EDTA, pH 8.0) and centrifuged again. Cell pellets were resuspended in 1 ml TES buffer, and 500 μl each was transferred to fresh 2 ml tubes. After two freeze-thaw cycles in liquid nitrogen and defrost at 60°C 5 mg ml^-1^ lysozyme (Carl Roth) and 0.5 μl RNase A (Thermo Scientific™, 0.1 μg ml^-1^ final) was added and incubated for 1 h at 37°C. Next, 2 μl ml^-1^ proteinase K (Carl Roth) and 2% SDS (ROTI^®^Stock 20% SDS, Carl Roth) were added and incubated overnight at 60°C. After incubation, 1 vol. (1:1 (v/v)) Phenol/Chloroform (ROTI^®^Phenol /Chloroform/Isoamyl alcohol, Carl Roth) was added and centrifuged at 12,000 g for 10 min, 4°C. The upper aqueous phase was transferred to a new tube and again 1 vol. 1:1 (v/v)) Phenol/chloroform (ROTI^®^Phenol/Chloroform/Isoamyl alcohol, Carl Roth) was added and centrifuged at 12,000 g, 10 min, 4°C. Samples were precipitated with 0.7% isopropanol (2-Propanol, ROTIPURAN^®^ ≥99%, p.a., Carl Roth) overnight at -20°C. Isopropanol was removed by centrifugation, samples were washed in 70% ethanol, and air dried for ∼1 h, and the DNA was solved in 30 μl TE buffer. Before the DNA was used, the samples were incubated at 4°C overnight.

### Strain construction

*Synechocystis kai* gene knockouts were generated by replacing the gene or gene cluster of interest with a kanamycin resistance cassette. DNA from the segregated mutants was isolated and used for the transformation of *Synechocystis* – substrain Kazusa (non-motile, derived from Uppsala University). For the transformation, 10 ml of bacterial culture were grown to an OD_750nm_ of 0.5 to 1, centrifuged at 2,000 g for 10 min, RT. The supernatant was discarded, and the pellet was resuspended in the remaining liquid and transferred to a new 1.5 ml tube. 20-50 μg ml^-1^ isolated DNA was added, and the sample was incubated for at least 30 min at 30° C, and subsequently transferred to a BG11 agar plate without antibiotics. After incubation at RT for at least 2 h, plates were placed in the Multitron Infors HT^®^ Incubator (see strains and growth conditions) for two to three days. For screening of desired colonies, 400 μl Kanamycin (1 mg ml^-1^) was applied to one side of the plate to form a gradient. For segregation, the kanamycin concentration was increased from 25 μg ml^-1^ to 50 μg ml^-1^, subsequently.

To generate the *glgC* (slr1176) knockout mutant, a fragment comprising the open reading frame for slr1176 plus 456 nt upstream and 843 nt downstream was amplified using the primers JS69 (5’ GGCATCAACGGCGTTGGAAA 3’) and JS70 (5’ GGCACCACTTCCACCGACTG 3’). The fragment was ligated into the cloning vector pJET1.2 (ThermoFisher). To obtain the kanamycin resistance cassette, the plasmid pUC-4K was digested with HincII. After purification, the kanamycin resistance cassette was inserted into the unique PsiI site of *glgC*. All restriction enzymes were purchased from New England Biolabs, USA. Transformation, selection on kanamycin-containing BG11 plates, segregation of independent clones, and verification by PCR analysis were performed as described in Eisenhut et al. 2006 (83).

### Growth experiments

Growth experiment pre-cultures were initiated from BG11 agar plates and propagated in 100 ml BG11 within 250 ml Erlenmeyer flasks for a minimum of five days. These cultures were then diluted to an OD_750nm_ of approximately 0.5 (Specord 200 Plus - Analytic Jena^©^), and grown until they reached an OD_750nm_ of approximately 2. When the desired OD_750nm_ was reached, the cultures were split and diluted to an OD_750nm_ of 0.8 and were placed on the Cell Growth Quantifier (CGQ, Aquila Biolabs, Scientific Bioprocessing (sbi) now) in an Infors Incubator for measurement of the backscatter signal at 730 nm. The CGQ-system consists of one base station, eight sensor plates with backscatter sensors for the flasks, and the CGQ live monitoring software.

### Temperature compensation

For the temperature compensation tests, we followed the previously outlined growth protocol with the modification of adjusting to the respective temperatures (25°C n=3, 30°C n=11, 35°C n=1, ‘n=’ denotes the number of independent experiments measured in our study) for *Synechococcus* and *Synechocystis*, which were cultured in parallel, at the respective temperatures. The data from repeated measurements of the same samples were then analyzed by calculating the mean from four periods of each measurement. Q_10_ values, were calculated from 25°C and 30°C and were validated with the 35°C measurement.

### Entrainment of the cells

Entrainment of the clock was tested with two different *Zeitgeber* signals, each of them applied with a 12 h delay to two different cultures. For the first approach, we mimicked a nutrition upshift (n=3) by diluting the culture with media: We inoculated two separated cultures in BG11 and let them grow in the incubator to an OD_750nm_ of ∼2. Then the first culture was diluted to an OD_750nm_ of 0.8. 12 h later, the second culture was diluted to an OD_750nm_ of 0.8 and the backscatter signal was monitored in the above described CGQ system. As a second approach, we entrained cultures with LD cycles. As before, two separate cultures were grown to an OD_750nm_ of ∼2 and then diluted to an OD_750nm_ of 0.8. Both cultures were exposed to LD cycles (12:12, LD, n=3) over 48 h, but the first dark phase was shifted by 12h. Following the entrainment, measurements were initiated in the CGQ System, where ‘n=’ represents the number of independent experiments conducted, each consistently including at least two technical replicates.

### Cell Analysis

In the selected experiment, one batch culture (700 ml, OD750nm of 0.8) of *Synechocystis* WT was divided into seven batch cultures (100 ml each), while *Synechocystis* Δ*kaiA1B1C1* and *Synechocystis* Δ*glgC* were each divided into four batch cultures (100 ml each). For the WT culture, seven time points were selected, with samples taken every 2 hours for a time frame of 12 hours, starting at 86 hours after inoculation. For the Δ*kaiA1B1C1* and Δ*glgC* cultures, four time points were selected, with samples taken every 4 hours within the same 12-hour time frame. At each time point, a sample was taken from a different culture, with each culture being sampled only once. For all samples, we collected 5 ml for measuring glycogen content, 5 ml for determining cell dry weight, 1 ml for optical density, and cell count measurement.

#### Glycogen measurement

To determine the glycogen content, 5 mL of cell culture were harvested into a pre-cooled (−20°C) reaction tube. After centrifugation at 20,000 g for 5 min at 4°C, the supernatant was immediately discarded, and the cell pellets were flash-frozen in liquid nitrogen and stored at −80°C until further processing. For glycogen extraction, the method of Ernst et al. 1984 (84) was modified as follows: Each cell pellet was resuspended in 4 ml KOH (30% w/v) and was incubated at 95°C for 2 h. For precipitation, 3 new sample tubes were filled with 400 μl per sample point, 1.2 ml of ice-cold ethanol was added, and the mixture was incubated overnight at -20°C. After centrifugation at 4°C for 10 min at 10,000 g, the pellet was washed once with 70% ethanol and again with 100% ethanol. Afterward, the pellets were dried with the Concentrator Plus speed-vac (Eppendorf) for 20 min at 60°C. Then the pellet was resuspended in 1 mL 100 mM sodium acetate (pH 4.5) supplemented with amyloglucosidase powder (Sigma-Aldrich, 10115) to a final concentration of 35 U/mL. For enzymatic digestion, samples were incubated at 60°C for 2 h. For the spectrometric glycogen determination, the Sucrose/D-Glucose Assay Kit from Megazyme (K-SUCGL) was applied according to the manufacturer’s specifications, but omitting the fructosidase reaction step and scaling down the total reaction volume to 850 μL according to Behle and Dietsch *et al*. 2022 (6). Absorbance at 510 nm was measured using a BMG Clariostar Photospectrometer.

#### Cell dry weight

To determine the cell dry weight (CDW), 5 ml of *Synechocystis* cultures were sampled onto a Petri dish of known weight. The Petri dishes with the respective samples were dried at 60°C for 24 h and weighed again after cooling. A Petri dish with 5 ml dried growth medium (BG11) served as a reference.

#### Optical density

All comparative measurements of optical density were performed in a Specord 200 Plus (Analytic Jena©) dual path spectrometer using BG11 as blank and reference. All samples were diluted 1:10 before each measurement due to their high optical density.

#### Cell count and size

To determine the cell count and cell size, 10 μl of the samples used for optical density and absorption spectra measurements were further diluted with 10 ml CASYton (OLS) to a final dilution of 1:10^4^. The cell count was measured with electric field cell counting (Schaerfe Systems, CASY Cell Counter model TTC). A capillary with a pore size of 45 μM was used for the respective measurements, and all measurements were performed with 200 μl in triplicates. The cell size was recorded in the entire diameter range of 0 to 30 μm, but only the data range of 0 to 10 μm was used for evaluation. The counted events in electric field cell counting are a mixture of live and dead cells, as well as cell debris and background signals from the growth medium. Only counts with a diameter between 1.5 and 5 μm were considered for the experiment (6). We conducted six replicates for *Synechocystis* WT and four replicates for the Δ*kaiABC*. However, we were only able to conduct two experiments for the Δ*glgc* strain, as the capillary frequently clogged during these experiments, likely due to excreted material, a phenomenon further discussed in the subsequent section.

### Data analysis and visualization

The boxplots extend from the lower to upper quartile values of the data, with a line marking the median. The whiskers extend from the box to demonstrate the range of the data, with outliers being represented outside of the whiskers. For violin plots, we presented the distribution of data and its probability density. The width of the ‘violin’ reflects the frequency of the data at different values. Data processing and visualization were done with Python 3.0. Built-in functions from “SciPy” and “NumPy” were used for independent t-tests and period determination.

The Q_10_ value compares the rate of biochemical processes or reactions over a temperature range of 10°C. A temperature-independent process has a Q_10_ value of 1. The temperature coefficient Q_10_-values for the evaluation of Temperature Compensation for frequency were calculated using the formula:

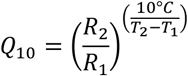

where the frequency (*R*_*x*_) is determined by calculating the period measured in the experiment and *T*_*x*_ is the corresponding temperature at which the experiment was conducted.

### Backscatter signal analysis

The aim is to detect periodic behavior in the time series data with a certain confidence. Measuring a parameter like a period can be challenging due to the non-stationarity and noisy nature of biological signals. Different Fourier-based and wavelet-based methods have been developed for detecting and analyzing circadian rhythmicity in biological data (85, 86). Unlike the Fourier transform, the wavelet transform has a high resolution in both the time and frequency domains. Wavelet allows studying the frequency components and time information of the data simultaneously. The time-averaged wavelet power spectrum of the time series data is used to determine the presence of a significant rhythm and the rhythm’s period. We analyzed the backscatter signals above a threshold of 330 [a.u.] and defined this point as t = 0) for comparability. For the mutant strains (Δ*kaiA1B1C1* and Δ*glgC*) no threshold was applied, because they behaved differently from the WTs and did not oscillate reliably after 330 [a.u.], but rather did not oscillate at all over the entire timescale. Experimental data were analyzed as described in Data preprocessing. Scripts are publicly available at github: https://github.com/rahilgholami/Circadian-rhythmicity.

### Data preprocessing

If, for a time series, statistical properties such as mean and variance change over time, the time series is non-stationary. We remove drifts in these statistical measures on time scales sufficiently longer than the expected 24 h period by fitting a linear regression model to the time series and then calculating the difference between observed values and predicted values from the model. Afterward, the time series is smoothed using the moving average (sliding window) as a part of the data preprocessing.

Periods were determined by measuring the time interval from peak to peak. For the detection of peaks the SciPy.signal.find_peaks” function was applied, which finds peaks inside a signal based on peak properties. This function takes a 1-D array and finds all local maxima by a simple comparison of neighboring values.

#### Wavelet transform

Suppose *x*_*n*_ is a discrete time series of *N* observations {*x*_*n*_, *n* = 0, …, *N*− 1} with a uniform time step *δt*. The continuous wavelet transform of the discrete time series *x*_*n*_is defined as

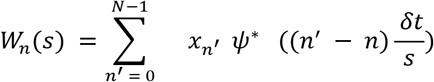

where *ψ*is the “wavelet function” and *s* is the wavelet scale. Larger scales stretch the wavelet function making it sensitive to lower frequencies in the signal. The wavelet power spectrum is defined as square of wavelet transform amplitude, ∣*W*_*n*_(*s*))∣^2^.

#### Significance level

Following Torrence and Compo 1998 (87) the statistical significance of the wavelet power can be assessed against a background power spectrum. A suitable background spectrum is either white noise or red noise.

Red noise can be modeled as a univariate lag-1 autoregressive process:

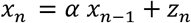

where *α* is being the assumed lag-1 autocorrelation and *z*_*n*_ is (Gaussian) white noise. The wavelet power spectrum of red noise is (87):

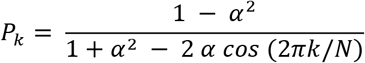

where *N*is being the number of data points and *π* is the frequency index. The mean background power spectrum significant at the 5% level is 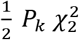, where 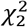is a chi-square distribution with 2 degrees of freedom.

## Acknowledgments

We thank Annegret Wilde for providing the *Synechocystis* Δ*kaiA1B1C1* mutant (26) for DNA isolation as well as her valuable comments on the manuscript. Thanks to Marion Eisenhut for assistance in reading the manuscript. We would like to acknowledge Lukas Becker and Nicolas Schmelling for their invaluable assistance with data analysis. Finally, we would like to especially thank Rainer Machné for continued discussions and critical reading of the manuscript.

Funded by the Deutsche Forschungsgemeinschaft (DFG, German Research Foundation) – Project ID 458090666 / CRC1535/1.

